## Supplementary material for "JAK-STAT pathway activation compromises nephrocyte function in a *Drosophila* high-fat diet model of chronic kidney disease": Figure supplement

### Supplemental documents

### SUPPLEMENTARY FIGURES

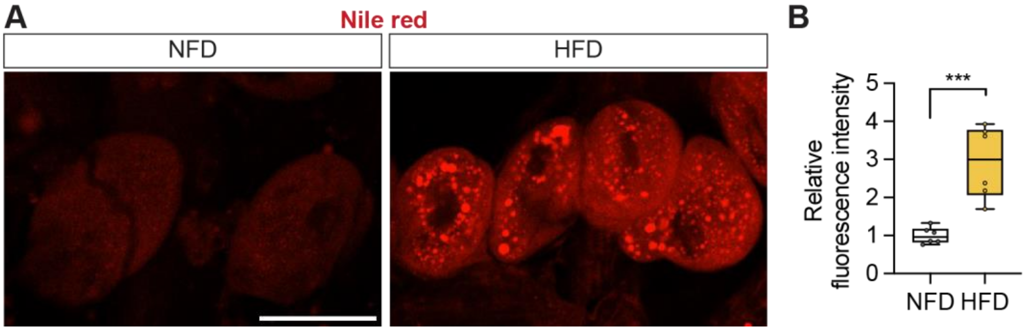


**Figure 1 - figure supplement 1. High-fat diet leads to lipid droplets accumulation in the nephrocytes**

**(A)** Nephrocytes from *Drosophila* $w^{1118}$ fed a regular diet (normal fat diet, NFD) or high-fat diet (HFD, NFD supplemented with 14% coconut oil). Nile red stains lipid droplets in red. Scale bar: 50 µm. **(B)** Quantitation of Sns-mRuby3 protein distribution (cytoplasmic vs membrane); middle line depicts the median and whiskers show minimum to maximum. Statistical analysis was performed with a two-tailed Student's t-test; ****, P<0.0001; n = 6 flies (7-day-old females).

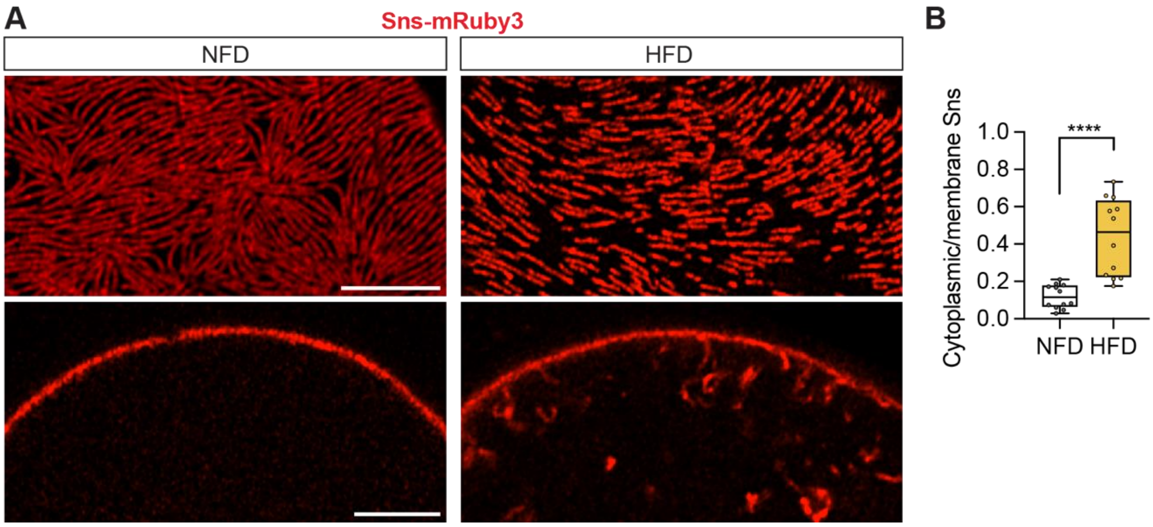


**Figure 2 - figure supplement 1. High-fat diet changes nephrocyte morphology**

**(A)** Nephrocytes from *Drosophila* (*sns-mRuby3*, 7-day-old females) fed a regular diet (normal fat diet, NFD) or high-fat diet (HFD, NFD supplemented with 14% coconut oil). Sns-mRuby3 is in green. Upper panels show cortical surface; Scale bar: 5 µm. Lower panels show subcortical regions; Scale bar: 5 µm. **(B)** Quantitation of Sns-mRuby3 protein distribution (cytoplasmic vs membrane); middle line depicts the median and whiskers show minimum to maximum. Statistical analysis was performed with a two-tailed Student's t-test; ****, P<0.0001; n = 12 nephrocytes (1 nephrocyte/fly) from 7-day-old female flies.

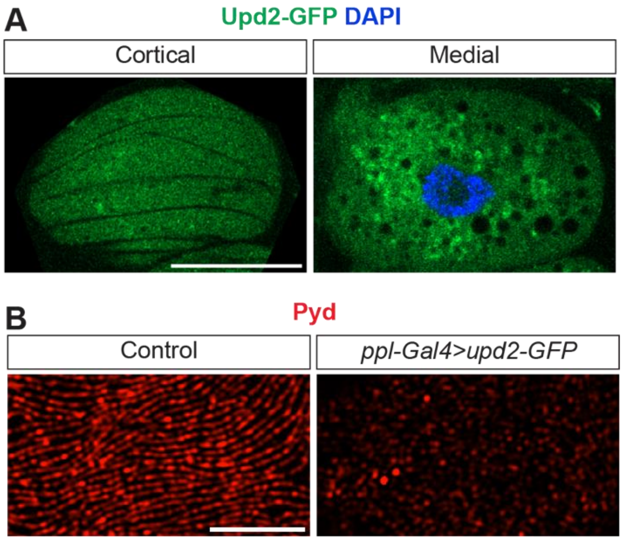


**Figure 5 - figure supplement 1. Upd2-GFP is secreted from the fat body and transported to the nephrocytes**

**(A)** Representative confocal images of nephrocytes. Genotype: *pp-Gal4>UAS-GFP*. GFP is shown in green. DAPI stains the nuclei in blue. Scale bar: 20 μm. **(B)** Representative confocal images of nephrocyte cortical regions. Anti-Pyd is shown in red. Scale bar: 4 μm.

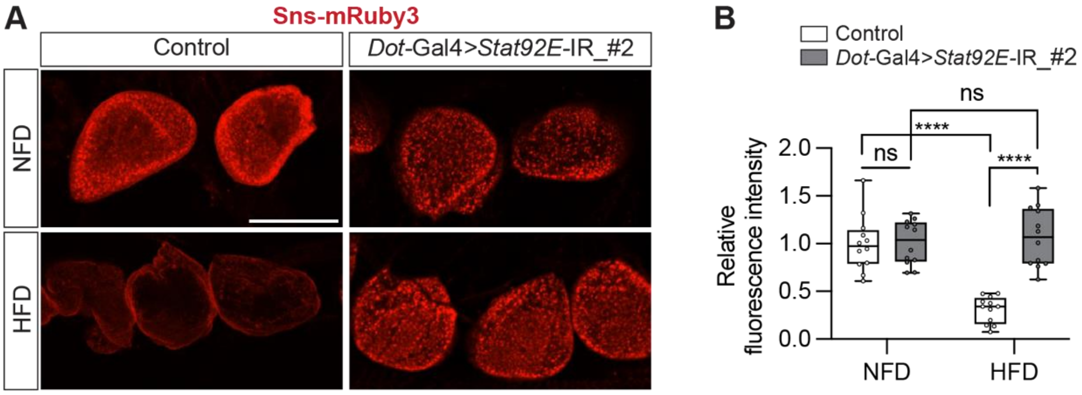


**Figure 6 - figure supplement 1. Stat92E depletion rescues HFD-caused nephrocyte functional decline.**

**(A)** Representative confocal images of nephrocytes from female adults that were fed on a regular diet (normal fat diet, NFD) or high-fat diet (NFD supplemented with 14% coconut oil, HFD) for 7 days. Genotype: Control (*Dot-Gal4-Gal4*/+); Stat92E depletion (*Dot-Gal4-Gal4*/+; *UAS-Stat92E-IR_*#2/+). Dextran is shown in red. Scale bar: 40 μm. **(B)** Box plot shows the quantitation of the relative fluorescence intensity of 10 kD dextran uptake based on images in (A); middle line depicts the median and whiskers show min to max. Statistical analysis was performed with a two-way ANOVA corrected with Tukey; ****, P<0.0001; ns, not significant; n = 12 flies.

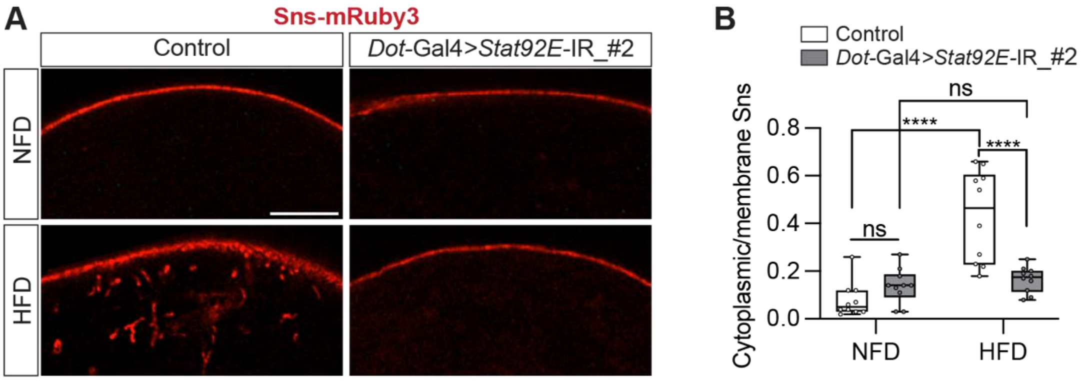


**Figure 6 - figure supplement 2. Stat92E depletion rescues HFD-caused Sns-mRuby3 distribution defects in the nephrocytes.**

**(A)** Representative confocal images of nephrocytes from female adults that were fed on a regular diet (normal fat diet, NFD) or high-fat diet (NFD supplemented with 14% coconut oil, HFD) for 7 days. Genotype: Control (*Dot-Gal4-Gal4, sns-mRuby3*/+); Stat92E depletion (*Dot-Gal4-Gal4, sns-mRuby3*/+; *UAS-Stat92E-IR_#2*). Sns-mRuby3 is shown in red. Scale bar: 5 μm. **(B)** Box plot shows the quantitation of Sns-mRuby3 protein distribution (cytoplasmic vs membrane) based on images in (A); middle line depicts the median and whiskers show min to max. Statistical analysis was performed with a two-way ANOVA corrected with Tukey; ****, P<0.0001; ns, not significant; n = 10 flies.

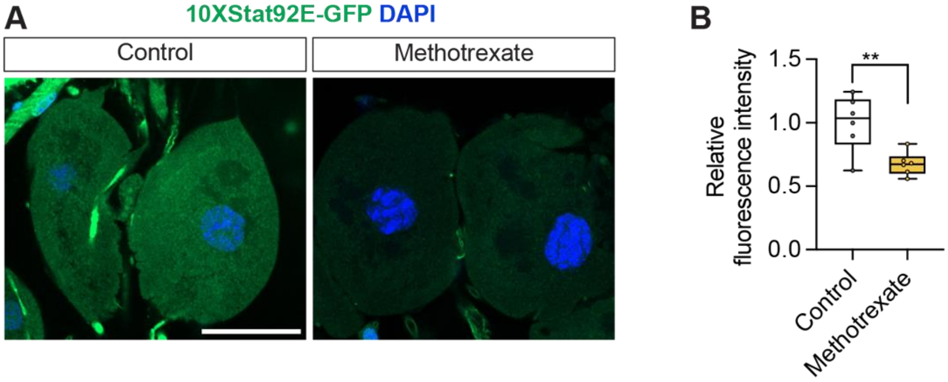


**Figure 7 - figure supplement 1. Methotrexate treatment inhibits JAK-STAT pathway activity**

**(A)** Representative confocal images of 7-day-old female adult nephrocytes (*10xStat92E*-GFP). Control, incubated in Schneider's Drosophila Medium (ex vivo for 60 min at room temperature); methotraxate, incubated in 10 μM methotrexate in Schneider's Drosophila Medium (ex vivo for 60 min at room temperature). *10xStat92E*-GFP in green fluorescence. DAPI staining in blue to visualize the nucleus. Scale bar: 20 μm. **(B)** Box plot shows the quantitation of the relative fluorescence intensity of *10xStat92E*-GFP based on the images in (A); middle line depicts the median and whiskers show Tukey. Statistical analysis was performed with a two-tailed t-test; **, P<0.01; n = 6 flies.

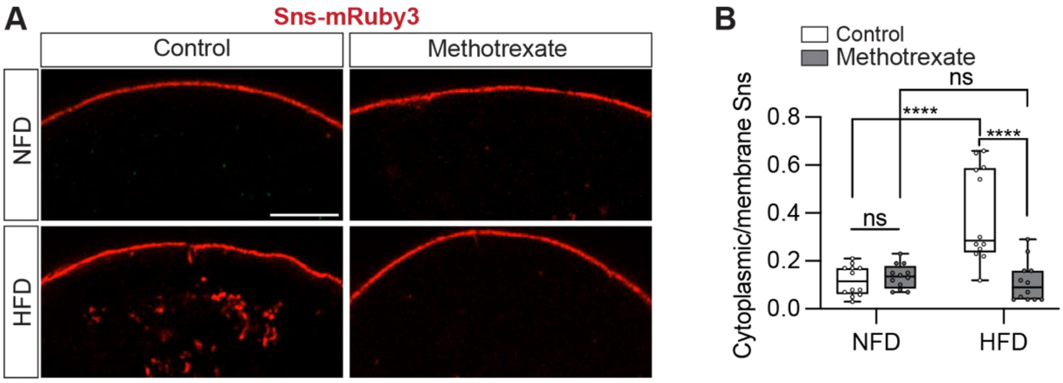


**Figure 7 - figure supplement 2. Methotrexate treatment restores Sns-mRuby3 distribution defects following a high-fat diet**

(**A**) Representative confocal images of nephrocytes from control *Drosophila* (*sns-mRuby3*; 7-day-old females) fed a regular diet (normal fat diet, NFD) or high-fat diet (NFD supplemented with 14% coconut oil, HFD), with or without methotrexate (10 µM; ex vivo 60 min) treatment. Sns-mRuby3 is in red. Scale bar: 5 µm. (**B**) Box plot shows the quantitation of Sns-mRuby3 protein distribution (cytoplasmic vs membrane) based on images in (A); middle line depicts the median and whiskers show minimum to maximum. Statistical analysis was performed by two-way ANOVA with Sidak correction; ****, $P<0.0001$; ns, not significant; n = 12 flies (7-day-old females).
